## supplementary figures for "A modified fluctuation assay reveals a natural mutator phenotype that drives mutation spectrum variation within *Saccharomyces cerevisiae*"

### Supplemental figures

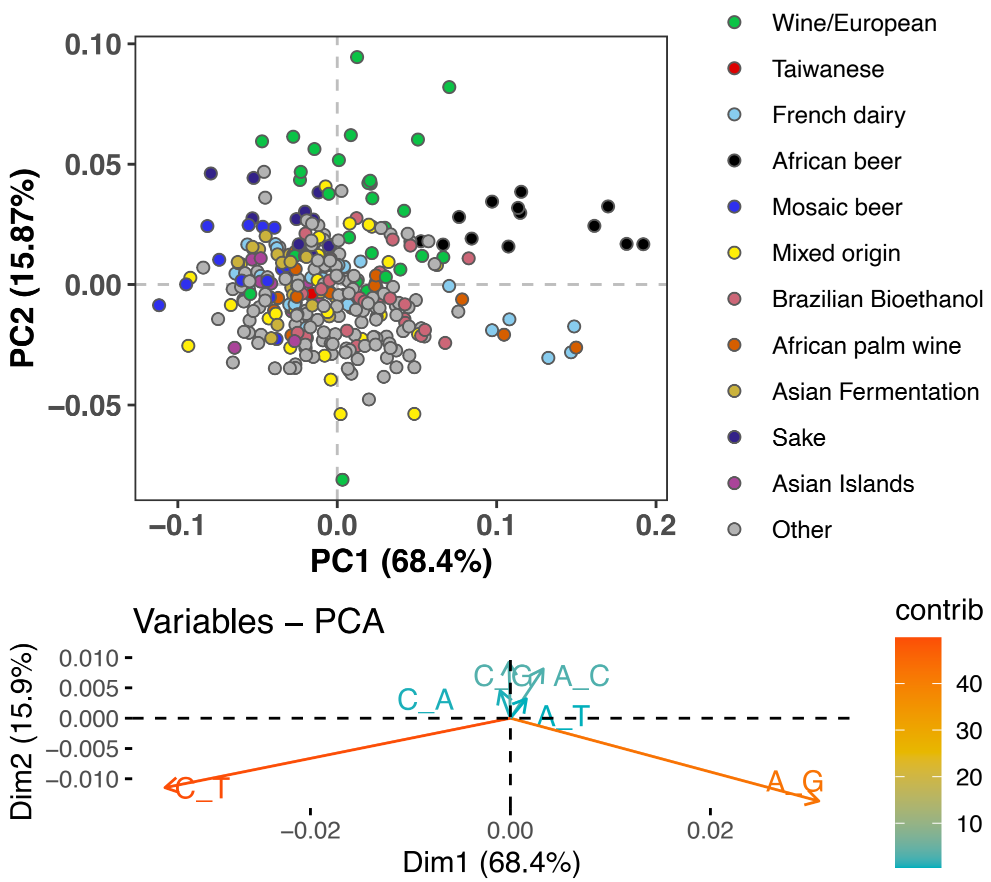


**Figure S1. Mutation spectrum PCA after subsampling to avoid overlap between lineages.** To eliminate any clustering of strains as a result of shared variation, we randomly assigned each mutation of frequency *k/n* to one of the *k* haplotypes carrying the derived allele, then computed each strain's spectrum based on these resampled mutation counts. The results still show similar clustering of outlier lineages to what is seen in Figure 1A without this subsampling, though with slightly more noise and dispersion.

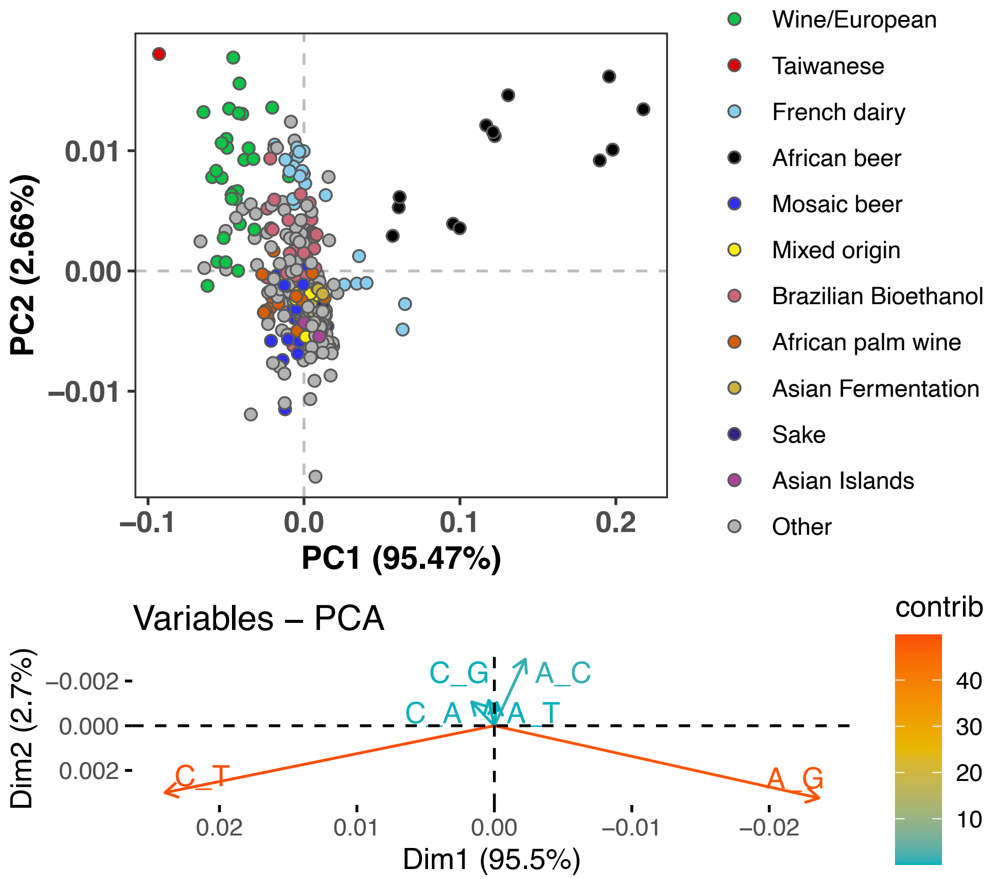


**Figure S2. PCA of synonymous variant mutation spectra**. To avoid the confounding effects of selection on nonsynonymous variants, we compute the spectrum of synonymous mutations present in each strain and normalize it by the spectrum of mutational opportunities for synonymous variants to arise. The resulting spectra display a similar PCA structure to what is seen in Figure 1A.

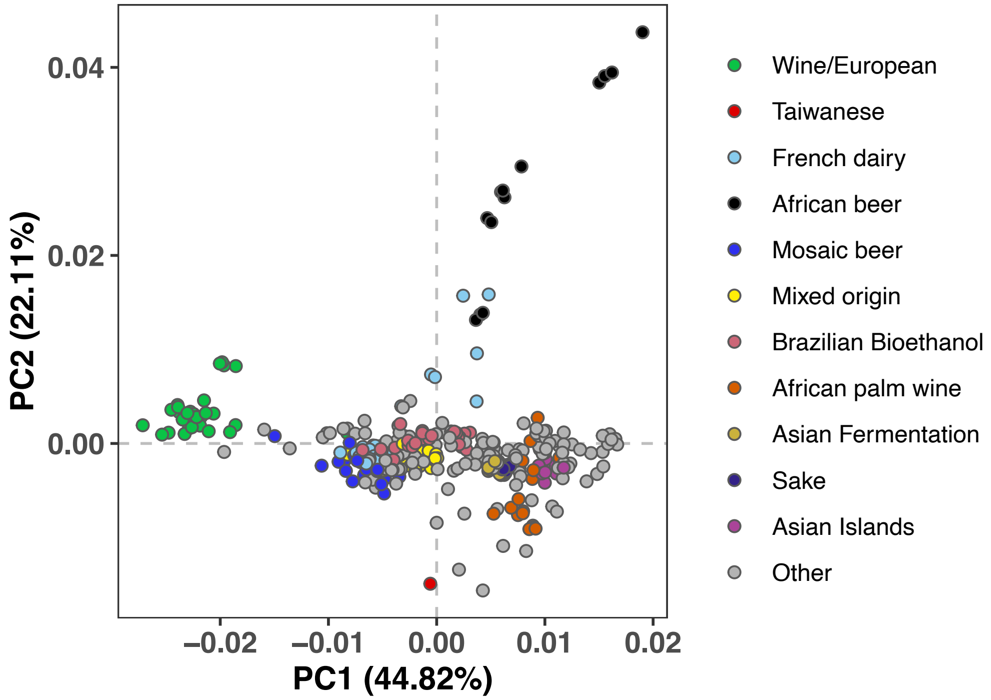


**Figure S3. Mutation PCA using all variants stratified by triplet context**. This PCA shows substructure among the 96-dimensional mutation spectra obtained by classifying all variants by the left and right adjacent base pairs as well as the ancestral and derived alleles. Clustering is qualitatively similar to what is observed in Figure 1A using 6-dimensional mutation spectra, classifying each variant by its ancestral and derived allele but no surrounding sequence context.

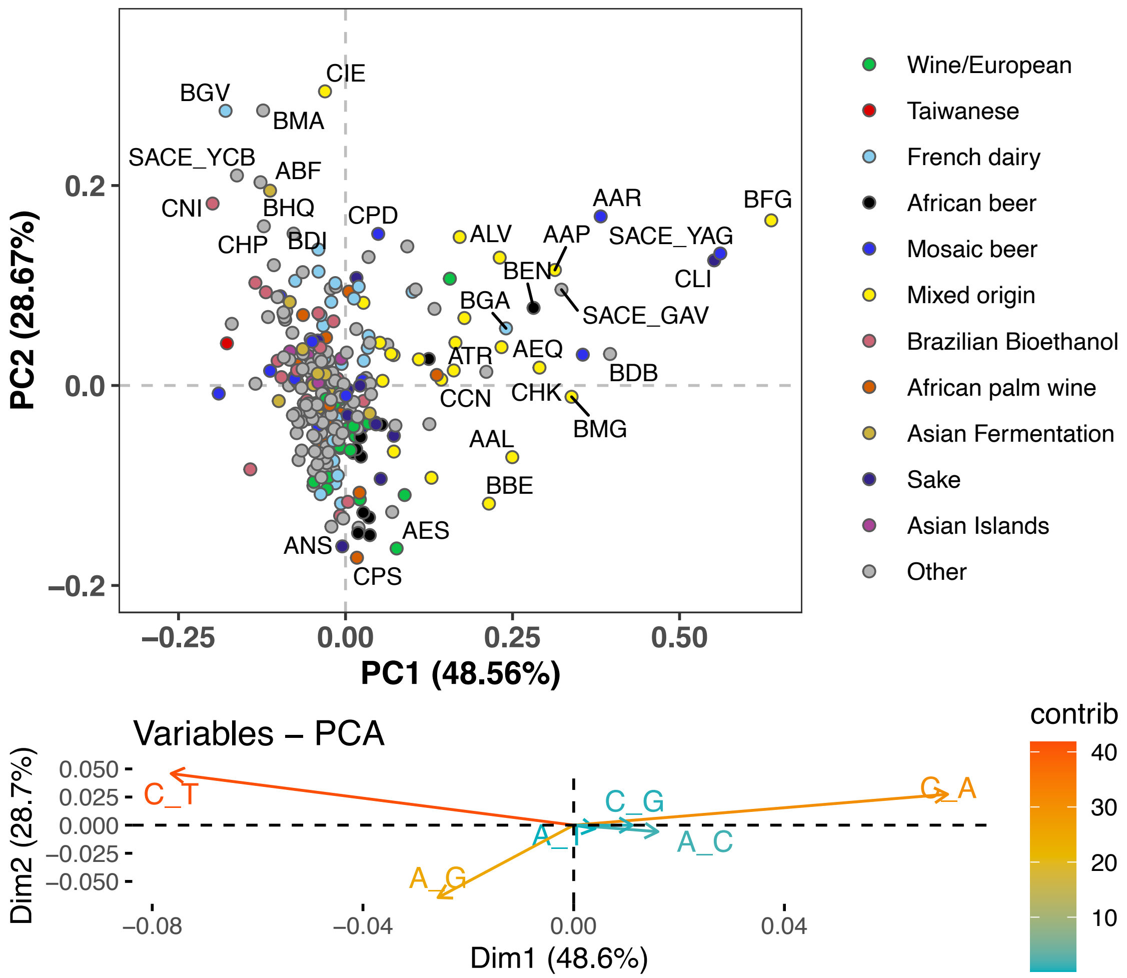


**Figure S4. PCA of singleton mutation spectra.** This PCA clusters mutation spectra that were computed using only singletons: variants present in the focal strain and no other strains. It shows a similar structure to Figure 1B, where spectra were computed from nonsingleton rare variants.

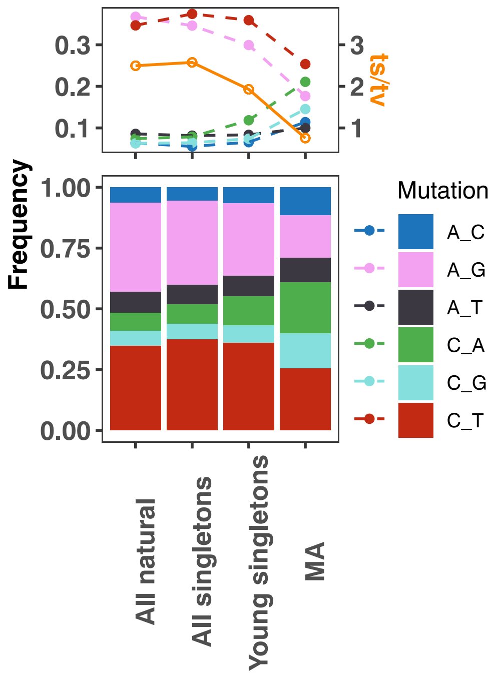


**Figure S5. Mutation spectrum comparison of natural variants versus *de novo* mutations from a previous mutation accumulation (MA) study.** From left to right are mutation spectra from 1) all natural variants, 2) all singletons, 3) young singletons (see Materials and Methods), and 4) MA experiments (Sharp et al. 2018, from haploid in RDH54+ backgrounds). These datasets are ordered such that mutations are expected to get younger from left to right. The top panel represents the frequency of each mutation type as a solid dot and the ts/tv ratio with an orange dot. The bottom panel shows the stacked frequency of each mutation type. Young singleton is defined similarly as in Zhu et al. 2017, and in this work, young singletons are defined by density of less than 0.0087 count/kb.

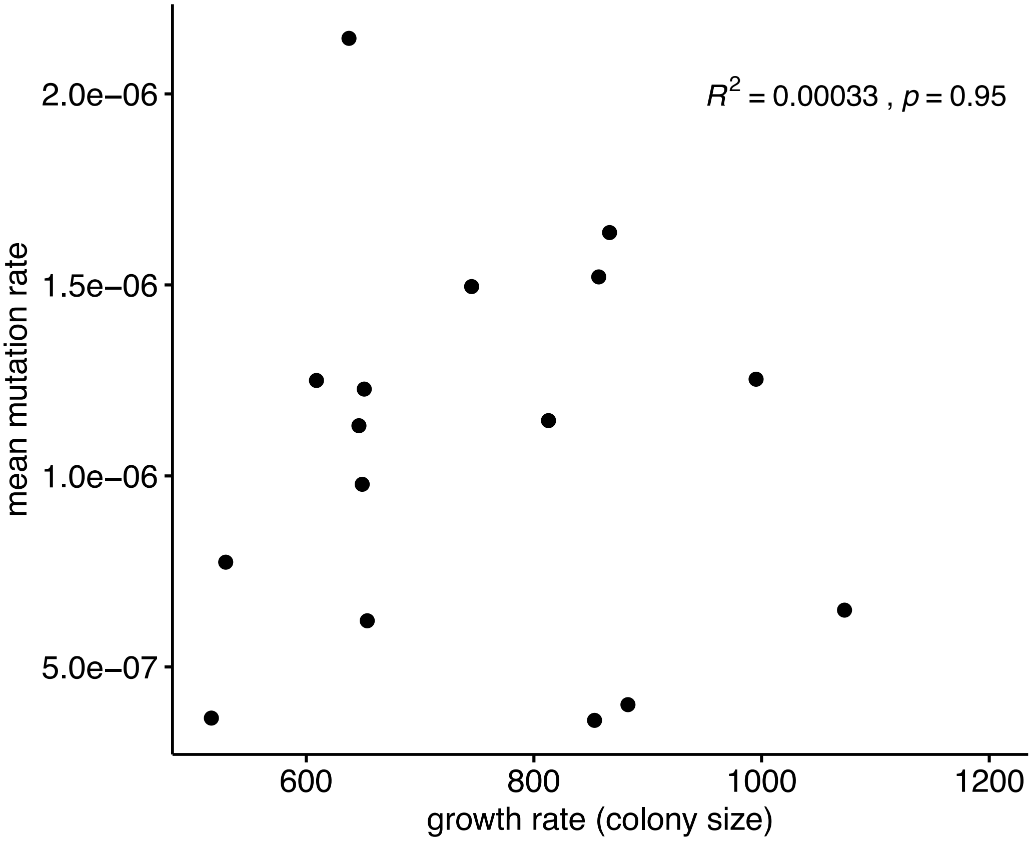


**Figure S6. Relationship between growth rate versus measured mean mutation rate.** Peter et al. 2018 measured growth rates from colony size from each of the 1011 yeast genomes. As shown, we see no evidence of a relationship between these variables for the strains included in our fluctuation study.

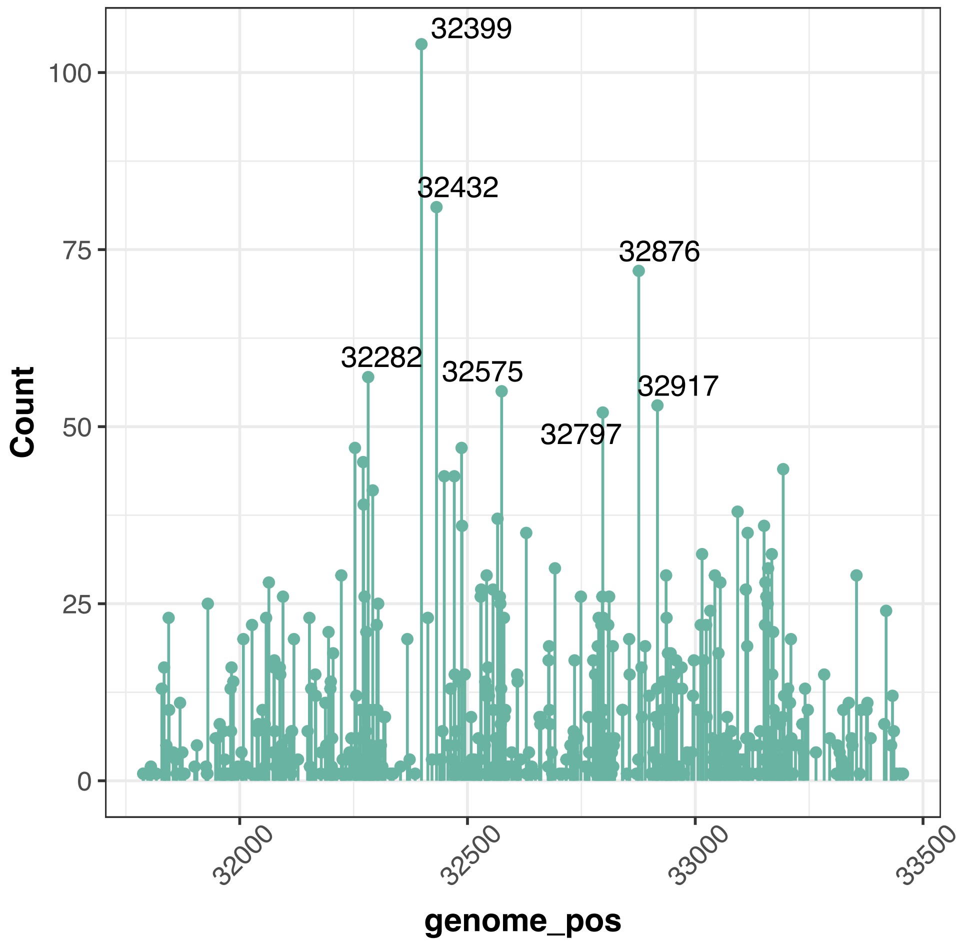


**Figure S7. Hotspots of *CAN1* mutation across different strain backgrounds (chr V).** Here, we plot the number of mutations observed at each genomic position across all fluctuation assays combined. Hotspots where 50 or more mutations were observed are labeled with their genomic positions.

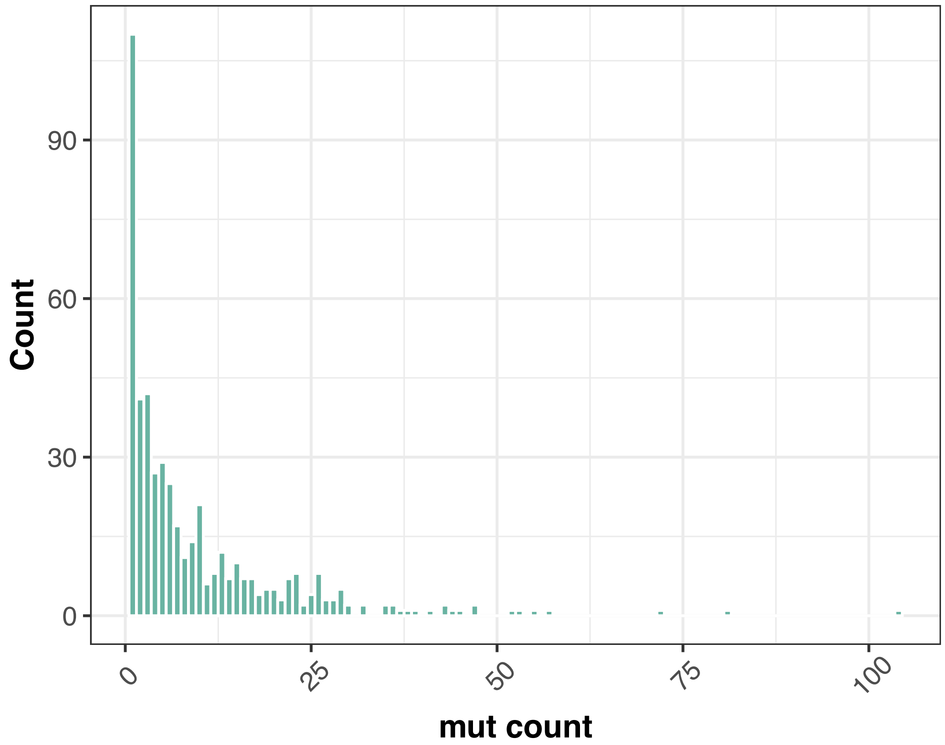


**Figure S8. Distribution of multiplicity of mutations observed at each mutated site in *CAN1*.** As summarized in this frequency spectrum, a plurality of mutations were observed just once, but some sites appear to be mutation hotspots that were found to be mutated in 50 or more independent fluctuation assays.

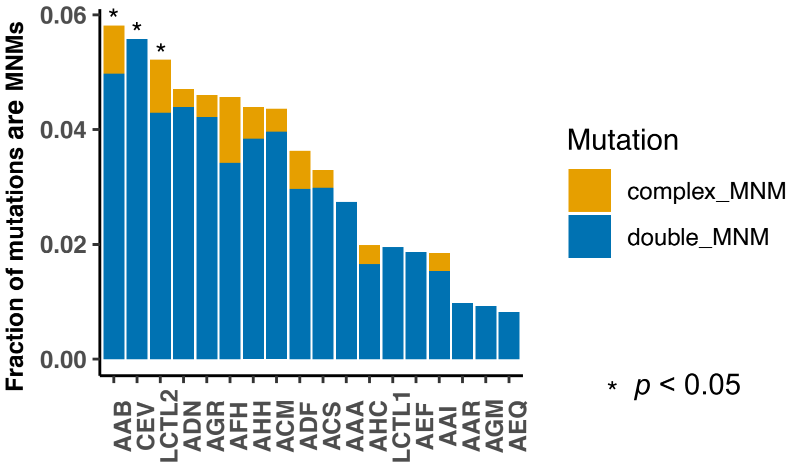


**Figure S9. Fraction of multinucleotide mutations (MNMs) in each strain.** A Chi-square test was performed on each strain to compare its ratio of MNM counts to single mutation counts to the ratio observed in the standard reference LCTL1 strain. Asterisks denote strains with significantly elevated MNM-to-SNP ratios ($p$<0.05).

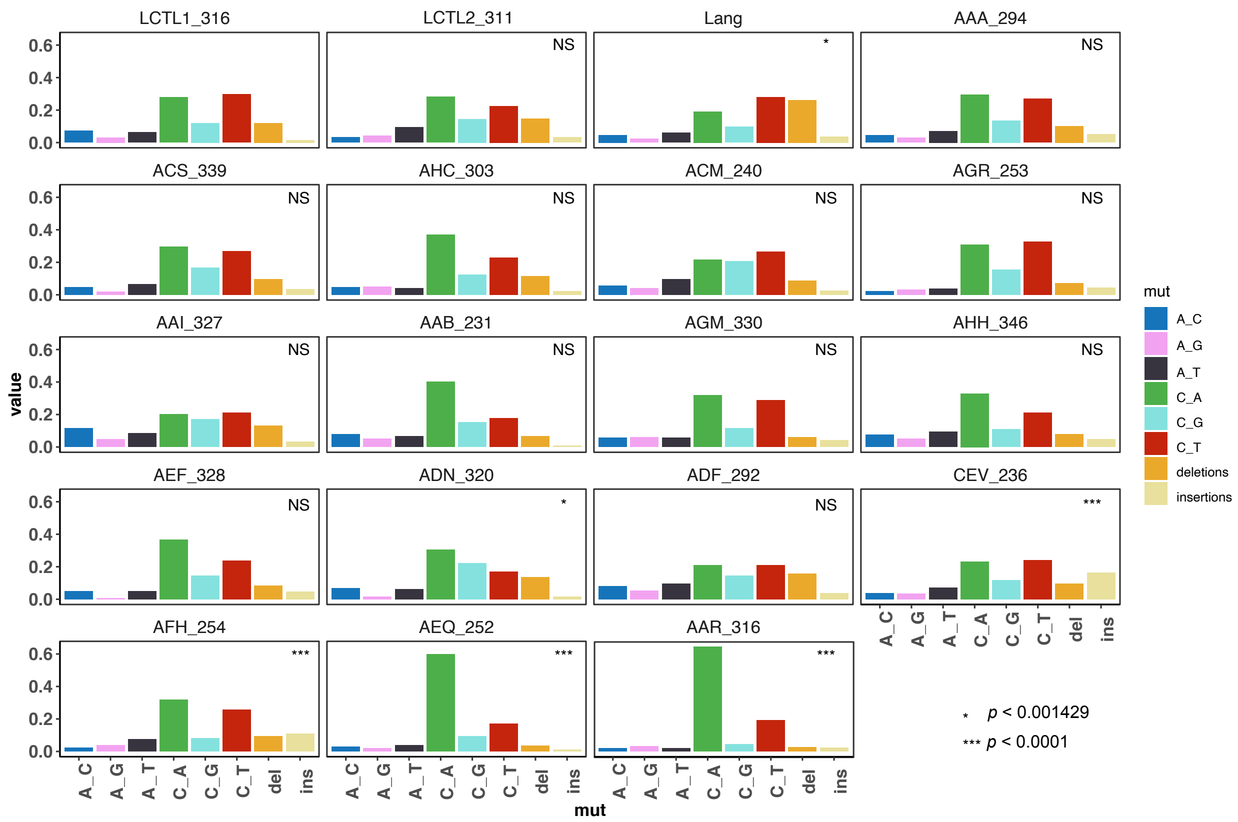


**Figure S10. *De novo* mutation spectra of all strains (single base-pair substitutions and single base-pair indels).** Each strain is marked by the strain name followed by the total number of mutants collected from that strain. We used a hypergeometric test to compare the mutation spectrum of each strain to that of the control LCTL1 strain, and those which differ significantly after Bonferoni correction are marked with an asterisk.

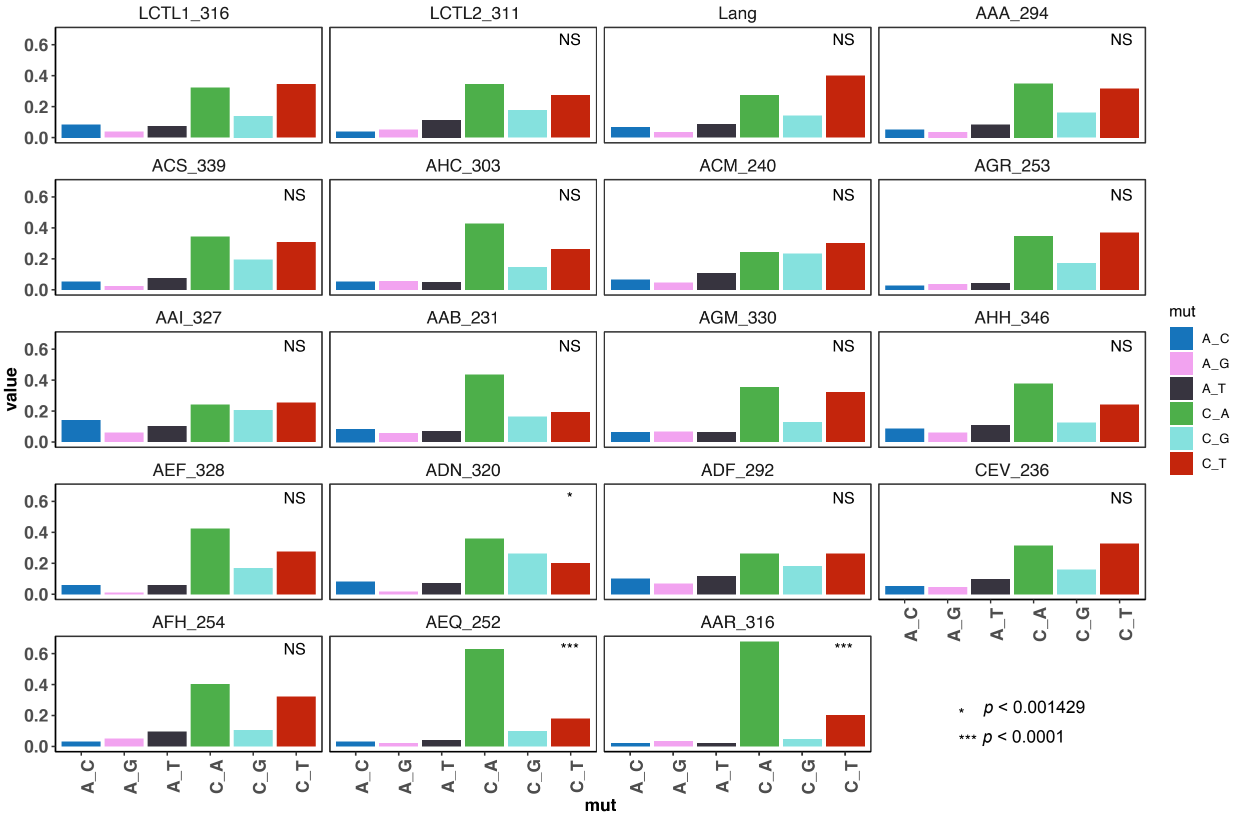


**Figure S11. D*e novo* mutation spectra of all strains (single base-pair substitutions only).** We used a hypergeometric test to compare each strain to the control LCTL1 strain, and those which differ significantly after Bonferoni correction are marked with an asterisk.

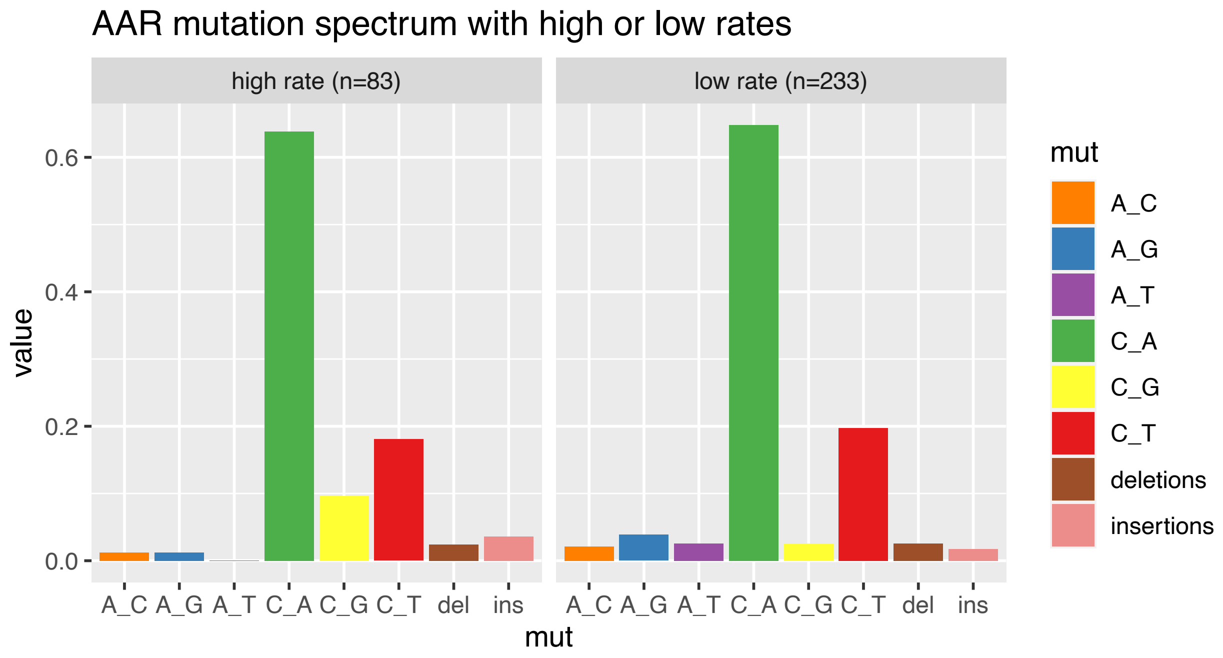


**Figure S12. Comparison of AAR mutation spectra from high versus low mutation rate batches.** As shown in Figure 3, we measured a bimodal distribution of mutation rates in the strain AAR. To investigate whether AAR pools with different measured mutation rates also have different mutation spectra, we classified each replicate as a high-rate or low-rate replicate based on whether the rate from the replicate was greater or less than 1.9e-6, then computed the mutation spectrum of each rate bin. The two spectra both exhibit the strain's distinctive enrichment of C>A mutations.

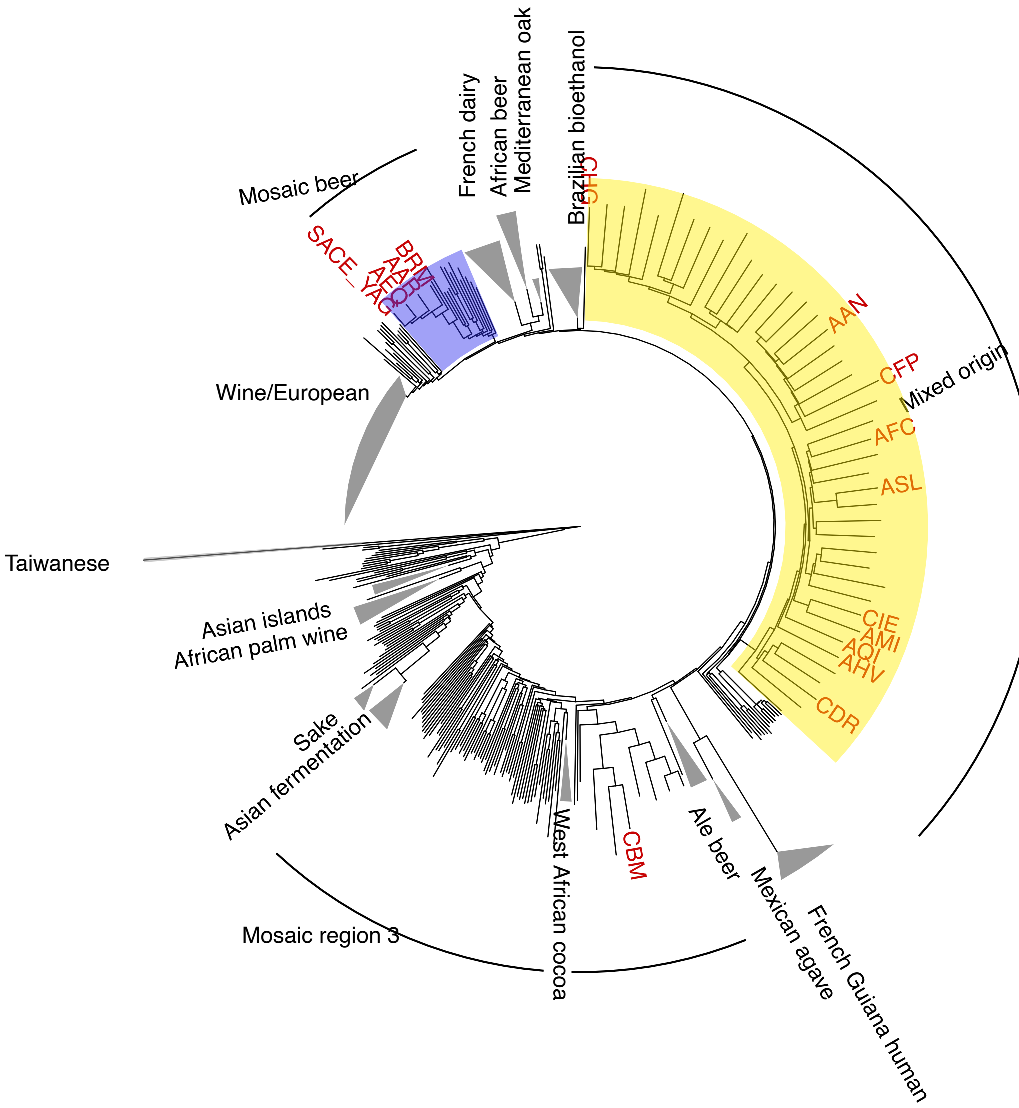


**Figure S13. Enrichment of C>A in rare polymorphisms in the 1011 collection (measured using empirical bootstrapping).** To further investigate the robustness of the enrichment of various lineages for rare C>A mutations, we compared their rare variant spectra to synthetic spectra obtained by subsampling rare variants from the 1011 collection. Each empirical spectrum was compared to 100 synthetic spectra created by subsampling variants to match the strain's total count of rare variants with global allele counts 2, 3, and 4. The strains highlighted here are the subset of the strains highlighted in Figure 5A that also appear significantly enriched for C>A by this bootstrapping approach, with an empirical $p$-value of less than 0.05.

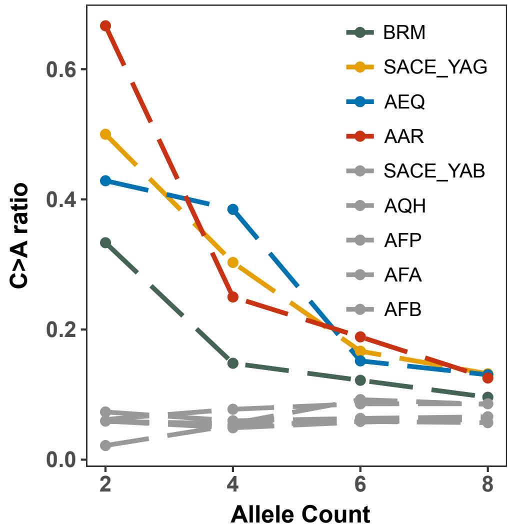


total mutation

| | | | | |
|---|---|---|---|---|
| AAR | 9 | 36 | 106 | 255 |
| AEQ | 7 | 26 | 112 | 230 |
| SACE_YAG | 6 | 33 | 108 | 250 |
| BRM | 12 | 54 | 123 | 271 |

**Figure S14. Enrichment of C>A mutations in rare natural variants from Mosaic beer strains that are closely related to AEQ and AAR from the 1011 collection.** C>A ratios in polymorphisms were calculated across allele count (AC) bins with cutoffs of 2, 4, 6, and 8. These allele counts are based on variation from the 1011 yeast genomes. They differ slightly from the counts in Figure 5A which include both the 1011 yeast genomes and the strain CBS1782.

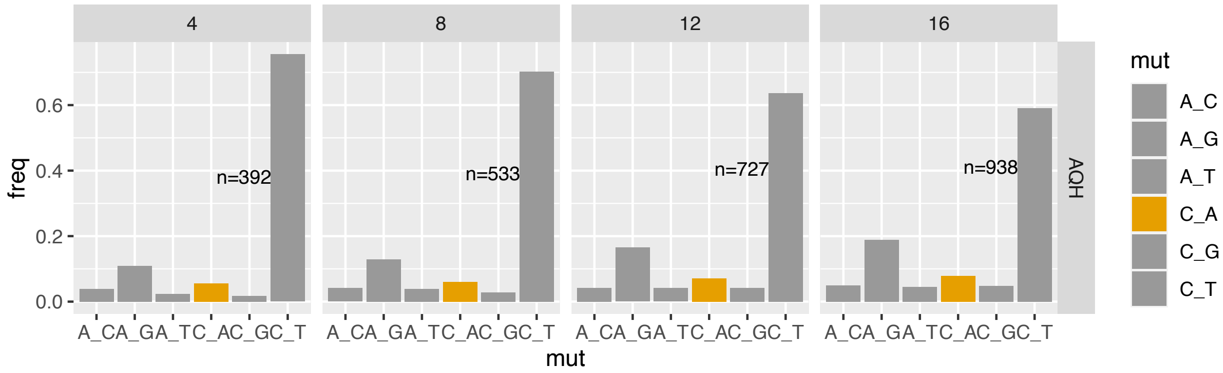


**Figure S15. Mutation spectra of rare natural polymorphisms in AQH stratified by minor allele count.** Each panel is labeled with its maximum allele count (AC) cutoff, meaning all variants with allele count less than or equal to the cutoff are included. Singletons are excluded.

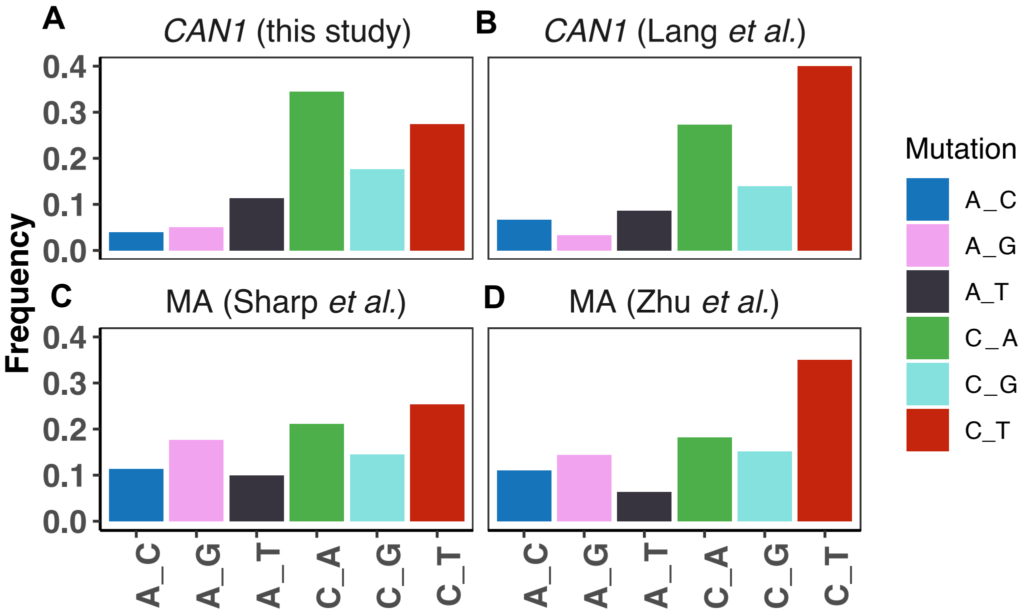


**Figure S16. Comparison of mutation spectra from *CAN1* reporter assays versus whole genome mutation accumulation (MA).** A. The *CAN1* mutation spectrum from the strain LCTL2 measured in this study (the same strain was previously used in Lang *et al*. 2008). B. The *CAN1* mutation spectrum of LCTL2 previously reported in Lang *et al*. 2008. C. A whole genome mutation spectrum from an MA study by Sharp *et al.* 2018, using haploid yeast from the RDH54+ strain. D. A whole-genome mutation spectrum of diploid MA study by Zhu *et al.* 2014.
